# Dual perspective proteomics infectome profiling discovers *Salmonella* type III secretion system effector functions in macrophages

**DOI:** 10.1101/2021.09.01.458519

**Authors:** J. Geddes-McAlister, A. Sukumaran, S.L. Vogt, J.L. Rowland, S.E. Woodward, B. Muselius, L. Gee, E.J. Roach, C.M. Khursigara, B. Raupach, B.B. Finlay, F. Meissner

## Abstract

Intracellular bacterial pathogens have evolved sophisticated infection strategies, including the release and secretion of virulence factors to interfere with host cell functions and to perturb immune responses. For *Salmonella enterica* serovar Typhimurium (*S*. Typhimurium), the type III secretion systems encoded on *Salmonella* pathogenicity islands (SPI) 1 and 2 mediates invasion of the bacterium into innate immune cells and regulates bacterial replication and survival within the hostile environment of the host, respectively. Here, we explore the temporal and strain-specific dual perspective response of both the host and pathogen during cellular infection via quantitative proteomics. We report time- and pathogenicity island-specific expression and secretion of infection-associated proteins (i.e., SL1344_1263, SL1344_3112, SL1344_1563, and YnhG) and regulated immune response proteins in macrophage, including Cd86, Cd40, Casp4, C3, IL-1α, and Cd69). Through intracellular macrophage and *in vivo* murine models of infection, we reveal a role in virulence for three of the bacterial proteins (SL1344_1263, SL1344_1563, and YnhG), defining their importance as novel T3SS effectors. We characterize the temporal intra- and extracellular production of the effectors and identify their interaction networks in host cells, representing inhibitory and stimulatory pathways mounted by invading bacterial pathogens.

**Author Summary:** The relationship between a host and pathogen is intricate, and regulation of the host immune response correlates with the progressive timing of infection and tailored responses to the pathogen. Relying on detection and quantification of protein-level changes using mass spectrometry-based proteomics, we explore the production of known and novel effectors by *Salmonella* during intracellular survival within macrophage. Our results portray a role for these effectors in bacterial virulence using an *in vivo* murine model of infection, and we define a dynamic network of interaction between the effectors and host proteins. These interactions reveal opportunity for drug design to disrupt modulation of the host by the invading bacterium as a new strategy for combatting infection. Our approach is dynamic and universal, with the power to alter therapeutic discovery against infectious diseases.

## Introduction

The enteric, pathogenic bacterium *Salmonella* causes a wide variety of diseases, ranging from gastroenteritis to systemic infections [1]. For this Gram-negative pathogen, two distinct type III secretion systems (T3SS) represent specialized virulence factors, which play an important role in delivering effector proteins to host cells. For *Salmonella enterica* serovar Typhimurium (*S*. Typhimurium), invasion of the host cell depends on bacterial proteins encoded in the chromosomal locus, *Salmonella* pathogenicity island 1 (SPI-1). The genes of SPI-1 encode for components of a T3SS, regulatory proteins, along with secreted effector proteins and their chaperones, which collectively establish the initial phase of infection [2]. In addition to inducing host cell death, SPI-1 mediates host cell invasion, autophagy, and inflammation [3–5]. Conversely, the *Salmonella* pathogenicity island 2 (SPI-2)-encoded T3SS permits replication of bacterial cells once inside macrophages by formation of a protective compartment, the *Salmonella* containing vacuole (SCV) [6–8]. Once inside the SCV, *Salmonella* is able to replicate and disseminate to systemic sites, including the spleen and liver, causing severe diseases (e.g., bacteremia) [9]. Currently, the increasing prevalence of global infection and multi-drug resistant *Salmonella* strains demands the discovery of new treatment options [10]. Understanding the mechanisms associated with *Salmonella* infection suggests innovative strategies for identifying novel therapeutics.

Previous work provides critical insight into the mechanisms of, and effectors associated with, the T3SS in *Salmonella* [6,9,11–14]. In addition, recent studies of the interaction between *Salmonella* and the host have defined effector-specific cellular reprogramming and options for drug discovery [15, 16]. The development of network resources (e.g., SalmoNet) for biochemical modeling, host-pathogen interaction studies, drug-discovery, experimental validation of novel interactions, and information to reveal new pathological mechanisms improves our understanding of *Salmonella* pathogenesis [17]. To uncover such information, mass spectrometry-based proteomics provides a robust, sensitive, and unbiased platform for profiling protein-level differences in complex biological samples [18]. Information about abundance, interactions, regulation, and post-translational modifications are defined, providing novel biological insight into bacterial pathogens and their relationship with the host [19, 20]. The majority of experiments focus on a single perspective (i.e., identification of pathogen or host proteins) given limitations of detection of pathogenic proteins in the presence of highly abundant host cells. For example, cellular proteome profiling of *Salmonella* from infected host epithelial cells focuses on the bacterial adaptive response to infection [21, 22]. Other studies use quantitative proteomics combined with high-throughput techniques for the characterization of host gene responses and host phosphoproteome dynamics following *Salmonella* infection by focusing on the impact of specific effectors [11, 23]. In addition, secretome profiling of *Salmonella* identifies a novel virulence factor and defined host epithelial cell response to infection, and proteomic profiling of *Salmonella*- induced filaments reveals critical information about bacterial survival mechanisms [24–27]. Moreover, interactions between the host and pathogen are explored using affinity purification and modern proximity labeling techniques (e.g., BioID), which provide valuable information into the complex network of interactions between a single effector and the host during infection [28, 29].

Building upon this previous knowledge and providing a universal platform to investigate the proteome of host-pathogen interactions from dual perspectives, we unbiasedly define the interaction between *Salmonella* and primary macrophages for both pathogenicity islands in a single experiment. In addition, our in-depth infectome profiling defines known and novel *Salmonella* effectors regulated by the pathogenicity islands and we uncover new temporal and strain-specific responses of the host to infection. Moreover, we characterize the novel effectors with *in vitro* and *in vivo* infection assays to reveal roles in virulence and visualize their production in host cells. Interactome profiling of the bacterial effectors reports unique points of biochemical contact within host cells, suggesting new opportunities to perturb such interactions as novel anti-virulence strategies for therapeutic intervention. Overall, this study is the first to report extensive protein-level dynamics during *Salmonella* infection simultaneously from global, host, and pathogen perspectives in consideration of both the SPI-1 and SPI-2 modes of action. Finally, our approach is universal for the discovery of novel infection-associated proteins and has applications with medical and agricultural relevance to combat microbial infections on a global scale.

## Materials and Methods

### Bacterial strains and growth conditions

*S*. Typhimurium SL1344 wild-type (WT) (streptomycin resistant), SL1344 Δ*spi1* (kanamycin resistant) by deletion of the pathogenicity island, and SL1344 *spi2-*defective (kanamycin resistant) by transposon insertion in *ssrA* by signature-tagged mutant (STM) construction, and LT2 were used for this study (strains generously gifted from A. Zychlinsky’s lab) [30, 31]. *Salmonella* strains were maintained on Luria broth (LB) plates with the respective antibiotics, followed by inoculation of LB broth (with antibiotics) with shaking (200 rpm) overnight (18 h) at 37°C. For *Salmonella in vitro* cultures, WT, Δ*spi1*, and *spi2-*defective were grown in LB (without antibiotic) to early stationary phase and collected for proteome analysis. The *in vitro* growth assay was performed in biological quadruplicate and the experiment was performed in triplicate.

### Bone marrow-derived macrophage collection and infection

Age-matched C57BL/6 female mice were euthanized by CO_2_ asphyxiation and femurs were removed. Femurs were cleaned, and marrow was removed in bone marrow-derived macrophage (BMDM)-Dulbecco’s modified eagle’s medium (DMEM) media (DMEM, 10% fetal bovine serum (FBS), 5% horse serum, 20% L929-conditioned media, 1% HEPES, 1% sodium pyruvate, 1% L-glutamine, 5% penicillin/streptomycin [pen/strep]). Cells were spun down and resuspended in BMDM-DMEM. Cells were grown for 7–10 days before use.

For infection, BMDMs were seeded in 6-well plates at 1×10^6^ cells/well in BMDM-DMEM media without pen/strep overnight at 37°C, 5% CO_2_. Bacterial cells were grown in LB broth to early stationary phase (same cultures used for *in vitro* growth described above), spun down, and resuspended in phosphate-buffered saline (PBS) and diluted in BMDM-DMEM media. Cells were infected at a multiplicity of infection (MOI) of 50:1 (bacteria:macrophage) for 30 min at 37°C, 5% CO_2_. Subsequently, cells were washed with PBS and incubated at 37°C, 5% CO_2_ in growth medium containing 100 µg/mL gentamicin for 1 h. Medium was replaced to decrease the gentamicin concentration to 10 µg/mL for later time points. Cultures were collected at 0, 0.5, 1, 2, and 6 hours post infection (hpi) (i.e., hours post infection calculated following: 30 min co-culture, 1 h gentamicin treatment, add fresh DMEM) for proteome analysis. The BMDM infection assay was performed using biological quadruplicates and the experiment was performed in duplicate.

### Proteomic sample preparation

Total proteome, secretome, and infectome profiling was performed as previously described with minor modifications [32, 33]. Cell pellets (bacterial and macrophage) were collected and washed with PBS, resuspended in 100 mM Tris-HCl, pH 8.5 with proteinase inhibitor capsule (PIC) and mechanically disrupted by probe sonication in an ice bath. Next, 2% (final) sodium dodecyl sulphate (SDS) was added, followed by reduction with 10 mM dithiothreitol (DTT) at 95°C for 10 min at 800 rpm, alkylation with 55 mM iodoacetamide (IAA) at room temperature in the dark, and acetone precipitation (80% final) at -20°C overnight. Precipitated proteins were pelleted, washed twice with 80% acetone and air-dried, prior to solubilization in 8 M Urea and 40 mM HEPES. LysC was added at 1:100 µg protein for 3 h at room temperature, followed by addition of 300 µl 50 mM ammonium bicarbonate (ABC) and 1:100 µg protein of trypsin overnight at room temperature. Protein concentration was determined by BSA tryptophan assay [34]. For secretome samples, the supernatant from bacterial cell pellets above was filtered (0.22 µm syringe filter tip), reduced with 10 mM DTT for 30 min at room temperature, alkylated for 20 min with 55 mM IAA in the dark, and then subjected to LysC and trypsin digestion overnight as above. Digestion was stopped by addition of 10% v/v trifluoroacetic acid (TFA) per sample and the acidified peptides were loaded onto Stop-And-Go extraction tips (StageTips) (containing three layers of C_18_) to desalt and purify according to the standard protocol [35]. Approx. 50 µg of sample was loaded onto StageTips and stored at 4°C until LC-MS/MS measurement.

### Mass spectrometry

Samples were eluted from StageTips with Buffer B (80% acetonitrile (ACN) and 0.5% acetic acid), dried with a SpeedVac concentrator, and peptides were resuspended in Buffer A (2% ACN and 0.1% TFA). Approx. 3-5 µg of protein was subjected to nanoflow liquid chromatography on an EASY-nLC system (Thermo Fisher Scientific, Bremen, Germany) on-line coupled to an Q Exactive HF quadrupole orbitrap mass spectrometer (Thermo Fisher Scientific). A 50 cm column with 75 μm inner diameter was used for the chromatography, in-house packed with 3 μm reversed-phase silica beads (ReproSil-Pur C_18_-AQ, Dr. Maisch GmbH, Germany). Peptides were separated and directly electrosprayed into the mass spectrometer using a linear gradient from 5% to 60% ACN in 0.5% acetic acid over 120 min (bacterial cellular proteome and secretome) or 180 min (infectome) at a constant flow of 300 nl/min. The linear gradient was followed by a washout with up to 95% ACN to clean the column. The QExactive HF was operated in a data-dependent mode, switching automatically between one full-scan and subsequent MS/MS scans of the fifteen most abundant peaks (Top15 method), with full-scans (*m*/*z* 300–1650) acquired in the Orbitrap analyzer with a resolution of 60,000 at 100 *m*/*z*.

### Mass spectrometry data analysis and statistics

Raw mass spectrometry files were analyzed using MaxQuant software (version 1.6.10.43) [36]. The derived peak list was searched with the built-in Andromeda search engine [37] against the reference *S.* Typhimurium SL1344 proteome (downloaded from Uniprot, June 30, 2020; 4,658 sequences) and *Mus musculus* proteome (downloaded from Uniprot, June 30, 2020; 55,462 sequences). The parameters were as follows: strict trypsin specificity was required with cleavage at the C-terminal after K or R, allowing for up to two missed cleavages. The minimum required peptide length was set to seven amino acids. Carbamidomethylation of cysteine was set as a fixed modification (57.021464 Da) and N-acetylation of proteins N termini (42.010565 Da) and oxidation of methionine (15.994915 Da) were set as variable modifications. Peptide spectral matching (PSM) and protein identifications were filtered using a target-decoy approach at a false discovery rate (FDR) of 1%. ‘Match between runs’ was enabled with a match time window of 0.7 min and an alignment time window of 20 min. ‘Split by taxonomy ID’ was enabled when two proteome FASTA files were loaded in the same experiment (e.g., *S.* Typhimurium and *M. musculus*). Relative label-free quantification (LFQ) of proteins was performed using the MaxLFQ algorithm integrated into MaxQuant using a minimum ratio count of 1, enabled FastLFQ option, LFQ minimum number of neighbors at 3, and the LFQ average number of neighbors at 6[38].

Statistical analysis of the MaxQuant-processed data was performed using the Perseus software environment (version 1.6.2.2) [39]. Hits to the reverse database, contaminants, and proteins only identified by site were removed. LFQ intensities were converted to a log scale (log_2_), and only those proteins present in triplicate within at least one sample set were used for further statistical analysis (valid-value filter of 3 in at least one group). Missing values were imputed from a normal distribution (downshift of 1.8 standard deviations and a width of 0.3 standard deviations). A Student’s *t*-test was performed to identify proteins with significant differential abundance (*p-*value ≤ 0.05) (S0 = 1) between samples employing a 5% permutation-based FDR filter [40]. A Principal Component Analysis (PCA) was performed, as well as Pearson correlation with hierarchical clustering by Euclidean distance to determine replicate reproducibility within Perseus. STRING database was reference for interaction mapping (https://STRING-db.org) [41].

### Mutant construction

Mutants were first constructed in the *S*. Typhimurium LT2 background using the lambda Red recombinase method [42]. LT2 was used to improve transformation efficiency and ensure correct subsequent gene knockout in the SL1344 strains. Briefly, oligos were designed for homologous recombination of each gene (i.e., SL1344_3112, SL1344_1263, SL1344_1563, and *ynhG*) and used to amplify the kanamycin resistant cassette from pKD4 as previously described [15]. The WT LT2 was transformed with pKD46 (i.e., arabinose inducible lambda Red recombinase) at 30°C, followed by transformation of WT pKD46 with PCR product and plating on LB+Kan^25^ plates. Colonies were cured (37°C, overnight) of the pKD46 plasmid and confirmed for resistance cassette insertion into the gene of interest by colony PCR. Next, to transform into *S*. Typhimurium SL1344, the kanamycin resistance cassette was amplified from the LT2 background and transformed into SL1344 using the methods described above. At least two independent mutants were generated for each gene and mutants were confirmed by colony PCR and sequencing of the insertion sites. All primer sequences used in this study are provided (Supp. Table 1).

### Cytotoxicity assays

Murine primary macrophages were seeded onto a 12-well plate at a density of 0.1 × 10^6^ cells per well and incubated overnight at 37°C. Before infection with the bacteria, the medium was replaced with fresh BMDM-DMEM media (without antibiotics). Infection of cells with *Salmonella* was done as described [43], by adding bacteria 50:1 MOI onto the cells followed by incubation at 37°C. After 30 min of incubation, gentamicin (100 μg/ml) was added to kill extracellular bacteria, followed by incubation in fresh BMDM-DMEM media with gentamicin (10 μg/ml) . At the indicated time points (1, 3, 6, and 18 hpi), culture supernatants were collected for analysis as the experimental release samples. Cytotoxicity was quantified colorimetrically with the CytoTox96 lactate dehydrogenase (LDH)-release kit (Promega). The percentage of cytotoxicity was calculated with the formula: 100 × (absorption of supernatant/absorption whole cell lysate). The cytotoxicity assay was performed in biological triplicate and the experiment was performed in duplicate.

### Macrophage association, invasion, and replication

For phenotyping *S*. Typhimurium association, invasion, and replication (AIR) in primary macrophages, we followed the protocol previously described [44]. Briefly, primary macrophages were seeded at 0.1 x 10^6^ in BMDM-DMEN media overnight at 37°C, 5% CO_2_ and co-cultured with *S*. Typhimurium strains WT, Δ*spi-1*, Δ*spi-2*, ΔSL1344_1563, ΔSL1344_1263, ΔSL1344_3112, and Δ*ynhG* as described above. ‘Association’ plates were collected after 30 min co-culture, macrophage cells were lysed with 1% Triton X-100 for 10 min and released associated-*Salmonella* were serially diluted in PBS and plated on LB (WT) or LB+Kan^25^ (mutants). After 1 h of incubation in gentamicin, the ‘invasion’ plate was collected and treated as above, and after an additional 22.5 h, the ‘replication’ plate was collected, and serial dilutions were plated. Agar plates were incubated overnight at 37°C and CFUs were counted. Bacterial counts were normalized to macrophage cell counts. Comparison of association, invasion, and replication of normalized CFUs by Student’s *t*-test compared each strain to the WT and reported statistical significance (*p-value* <0.05). The experiment was performed in biological triplicate and repeated twice.

### Murine competitive index assays

All animal experiments were performed in accordance with the guidelines of the Canadian Council on Animal Care and the University of British Columbia (UBC) Animal Care Committee (certificate A17-0228). Mice were ordered from Jackson Laboratory (Bar Harbor, ME) and maintained in a specific pathogen-free facility at UBC. Ten six-week-old female C57BL/6 mice were orally gavaged with 100 µl PBS containing a total of 3×10^6^ or 3×10^7^ CFU of *S.* Typhimurium (in an approximate ratio of WT to mutant SL1344 of 1:1 as confirmed by retrospective plating of the inoculum). Mice were euthanized 3 days post-infection and tissue samples were collected for bacterial enumeration. Tissues were homogenized in PBS using a FastPrep-24 homogenizer (MP Biomedicals), serially diluted in PBS, and plated on LB+Str^100^ and LB+Str^100^+Kan^50^ for colony counts. Competitive index was calculated as the proportion of mutant (Kan^R^) *S.* Typhimurium in the tissue sample divided by the proportion of mutant *S.* Typhimurium in the inoculum. To ensure accurate calculations, competitive index was only determined for those samples having at least 50 colonies on LB+Str^100^.

### FLAG-tag proteins

Plasmids encoding C-terminally 3×FLAG-tagged versions of YnhG, SL1344_1263, and SL1344_1563 were generated by PCR amplification of the target gene from WT SL1344 genomic DNA. Each gene was expressed under the control of its native promoter; sequence encoding the 3×FLAG antigen was engineered into the reverse primer. PCR products were restriction digested and ligated into pACYC184 using standard techniques. The sequence of all inserts was verified by Sanger sequencing (UBC Sequencing + Bioinformatics Consortium).

### Western blots

Whole cell extracts and culture supernatants from *S.* Typhimurium WT and FLAG-tagged strains YnhG, SL1344_1263, and SL1344_1563 were fractionated by SDS-PAGE and transferred to a polyvinylidene difluoride membrane using a transfer apparatus according to the manufacturer’s protocols (Bio-Rad). After incubation with 3% non-fat milk in 1X TBS (50 mM Tris, 150 mM NaCl, pH 7.5) overnight at 4°C, the membrane was washed five times with TBST (1X TBS, 0.05% Tween-20), and incubated with ANTI-FLAG M2 antibody (Sigma-Aldrich) for 1 h. Membranes were washed three times for 5 min and incubated with 1:3000 dilution of horseradish peroxidase-conjugated Goat anti-mouse IgG Fc secondary antibody (Invitrogen) for 1 h. Blots were washed with TBST three times and developed using an alkaline phosphate (AP) substrate buffer (1 M Tris, pH 9.5, 5 M NaCl, 1 M MgCl_2_, 50 mg/ml 5-bromo-4-chloro-3-indolyl phosphate, and 5% nitrotetrazolium blue chloride). Experiment was performed in biological and technical triplicates.

### Immunofluorescence and imaging

For bacteria-only immunostaining, cultures were set overnight in LB broth prior to sub- culturing. At select times, bacterial cultures were collected and fixed in 80% methanol. Protocol for immunostaining was adapted from a previously described method [45]. Briefly, fixed cells were plated onto a cell slide coated with 0.1% poly-L-lysine (Sigma-Aldrich). Plated cells were incubated with buffer (25 mM Tris-HCl pH 7.6, 10 mM EDTA and 2 mg/mL lysozyme). Slides were submerged in 99% methanol, followed by 100% acetone prior to blocking with 10% goat serum in PBS. Antibodies were diluted in 1% BSA in PBS and incubated for 1 h. Cells were incubated with monoclonal ANTI-FLAG M2 antibody (Sigma-Aldrich) diluted to 1:100 followed by incubation with Anti-mouse Alexa Fluor 488 (AF488; Invitrogen) diluted to 1:200. Cells were counterstained with DAPI (10 µg/ml) for 1 min. Between incubation steps, slides were washed three times with PBS. A drop of *SlowFade^TM^* Gold antifade (Life Technologies) was added prior to mounting coverslips.

For infection immunostaining, C57BL/6 macrophages were maintained as described above. To minimize autofluorescence, cells were grown in FluoroBrite DMEM media (ThermoFisher Scientific) supplemented with 10% FBS 1 d prior to infection and during the infection protocol as described above. Samples were collected at 1 and 6 hpi and fixed overnight in 4% paraformaldehyde in PBS. Protocol for immunostaining was adapted from previously described methods and as outlined above [45, 46]. Cells were incubated with monoclonal Anti-FLAG antibody (Sigma-Aldrich) diluted to 1:100 followed by incubation with Alexa Fluor 488 diluted to 1:200. Cells were counterstained with DAPI (10 µg/ml) for 1 min. Between incubation steps, slides were washed three times with PBS. A drop of *SlowFade^TM^* Gold antifade (Life Technologies) was added prior to mounting coverslips.

Slides were imaged using a Leica DM5500B microscope, equipped with a Hamamatsu 3CCD digital camera operated through Volocity software ver. 6.3 (Quorum Technologies). For imaging macrophage samples, a fixed exposure of 500 ms was used to detect Alexa Fluor 488 bound to bacterial cells to account for the autofluorescence.

### Immunoprecipitation

FLAG-tagged *S*. Typhimurium strains and WT (as a negative control) were used to infect C57BL/6 macrophages as outlined above. At 1 and 6 hpi, cells were washed twice with wash buffer 1 (150 mM NaCl, 50 mM Tris, pH 7.5, 0.25% NP-40) and lysed with lysis buffer (150 mM NaCl, 50 mM Tris, pH 7.5, 5% glycerol, 1% NP-40, 1 mM MgCl_2_, and a PIC (Roche)). Lysates were diluted 1:1 with wash buffer 1 and incubated with Anti-DYKDDDDK Magnetic Agarose beads (Pierce) for 80 min with end-over-end rotation at 4°C. Beads were washed twice with wash buffer 1 and four times with wash buffer 2 (150 mM NaCl, 50 mM Tris, pH 7.5) followed by on-bead digestion as previously described [47]. Digested samples were purified by STAGE-tip and subjected to the LC-MS/MS protocol outlined above.

## Results

We devised a mass spectrometry-based proteomics approach to profile the abundance of bacterial proteins during laboratory growth (i.e., cell culture medium) combined with the infectome from dual perspectives (i.e., host and pathogen) in a single mass spectrometry measurement over the time course of infection (Fig. 1A). Our approach enables detection and tracking of both SPI-1- and SPI-2-associated bacterial proteins during infection of primary macrophages from invasion to replication, as well as the identification and characterization of novel infection-associated proteins.

**Fig 1.**
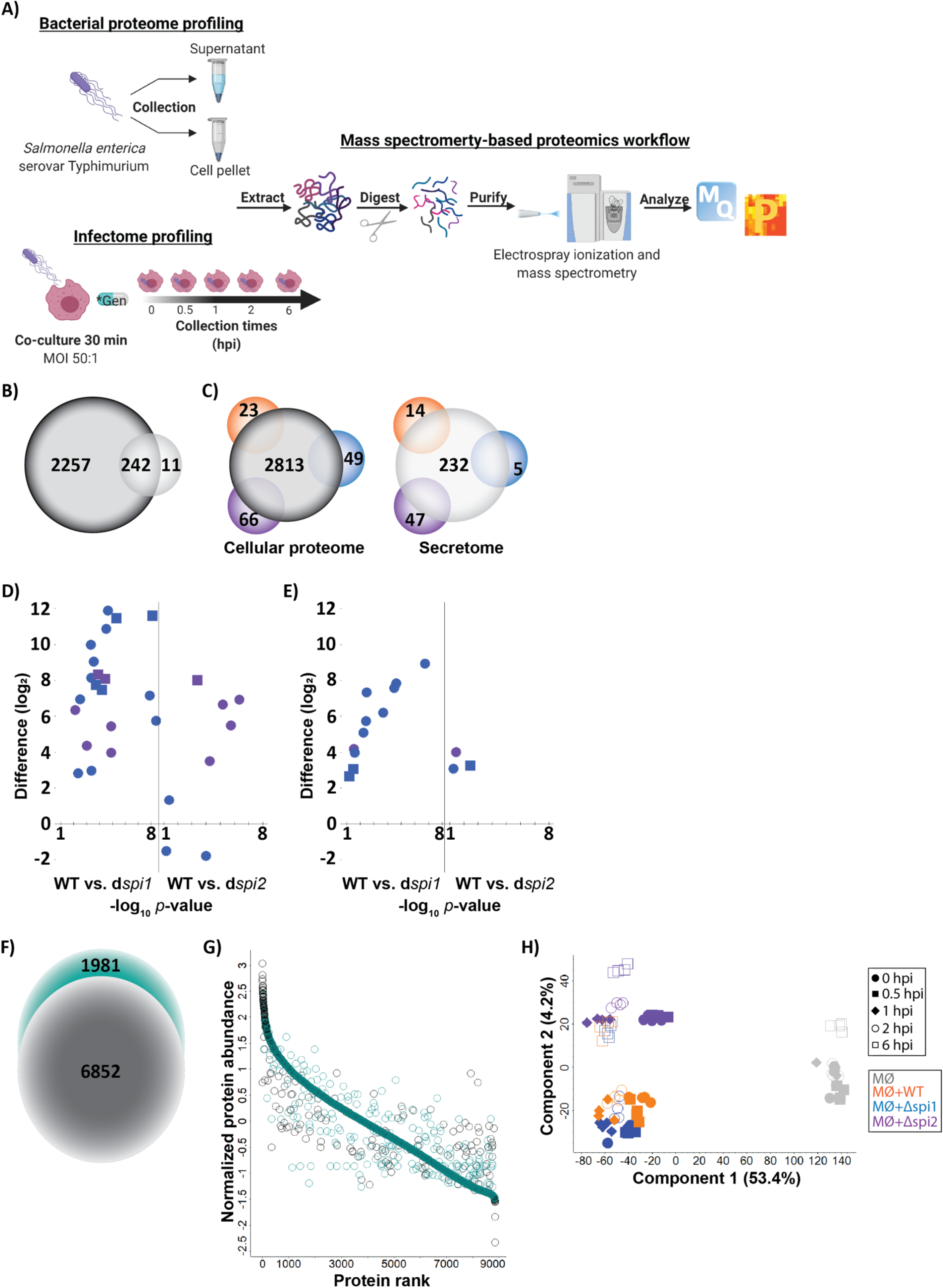
Mass spectrometry-based proteomics profiling of *S.* Typhimurium cellular infection. (A) *S.* Typhimurium cellular proteome (cell pellets) and secretome (supernatant) samples were collected and extracted followed by purification, electrospray ionization (ES), and liquid chromatography tandem mass spectrometry (LC-MS/MS). Data processing, analysis, and visualization performed with MaxQuant and Perseus platforms. Infectome timeline, collection, and processing of *S*. Typhimurium samples co-cultured with primary macrophages prior to mass spectrometry. Mass spectrometry-based proteomics experiments performed in biological quadruplicate and technical duplicate. *Gen = gentamicin treatment for 1 h following co-culture. (B) Venn diagram of proteins uniquely identified in the cellular proteome (2,257; dark grey), secretome (11; light grey) of *S*. Typhimurium under *in vitro* growth conditions, and proteins commonly found between the sample sets (242). (C) Venn diagram of a core bacterial cellular proteome (2,813; dark grey) and strain-specific proteins in WT (23; orange), Δ*spi1* (49; blue), and *spi2-*defective (66; purple); core bacterial secretome (232; light grey) and strain-specific proteins in WT (14; orange), Δ*spi1* (5; blue), and *spi2-*defective (47; purple). (D) Abundance of SPI-1- (blue) and SPI-2 (purple)-encoded or effector proteins in cellular proteome compared to WT. (E) Abundance of SP1-1- (blue) and SPI-2 (purple)-encoded or effector proteins in secretome compared to WT. Encoded = circles; effectors = squares. (F) Number of unique protein groups identified from host (6,852; grey) and *S.* Typhimurium proteins (1,981; green) across all time points (0, 0.5, 1, 2, 6 hpi) and bacterial strains (WT, Δ*spi1, spi2-*defective). (G) Dynamic range coverage (>5 orders of magnitude) of host (dark grey) and bacterial proteins (green). (H) Principal component analysis of infectome.

### Defining pathogenicity island-associated proteins *in vitro*

First, we analysed proteins produced in the cellular proteome and secretome across the three bacterial strains (Fig. 1B). From the cell pellet, we observed 2,499 bacterial proteins after filtering for valid values (representing 54% of the gene-encoding regions) and 11 uniquely secreted proteins, including proteins associated with the T3SS (*prgI*, *orgC*, *invJ*, *prgJ*, *spaO*), bacteriophage (SL1344_1940, SL1344_1941, SL1344_1953, SL1344_1958), and flagella (*fliK*). Among the *Salmonella* strains, we identified a core cellular proteome consisting of 2,813 proteins with 23, 49, and 66 proteins unique to WT, Δ*spi1*, and *spi2-*defective strains, respectively (Fig. 1C). Second, for the secretome, we observed consistent identification of 232 proteins with 14, 5, and 47 proteins unique to the WT, Δ*spi1*, and *spi2-*defective strains, respectively (Fig. 1C). Importantly, many of the unique proteins identified have roles as effectors or encoded genes for the respective pathogenicity islands, as evident in profiling differences in abundance of SPI-1- and SPI-2- associated proteins relative to WT in the cellular proteome (Fig. 1D) and secretome (Fig. 1E). As anticipated, these data show increased production of SPI-1- and SPI-2-associated proteins in the WT strain relative to the mutant strains, along with potential cross-regulation between the pathogenicity islands in the absence of *spi1*, possibly attributed to transcriptional regulation by *hilD*, as previously reported [48]. Notably, fewer SPI-2-associated effectors were detected in the secretome likely due to the selected growth condition (i.e., rich medium).

### Global profiling demonstrates depth and dynamic range of the infectome

To assess the depth of proteome coverage for both the host and pathogen in a single-run mass spectrometry experiment, we determined 78% were host proteins (6,852 host/8,833 total proteins) and 22% were bacterial proteins (1,981 bacterial/8,833 total proteins) (Fig. 1F). Notably, this represents detection of almost 80% of the total measurable bacterial proteome in the presence of the host. We observed a dynamic range spanning >5 orders of magnitude with distribution of both host and bacterial proteins (Fig. 1G). Finally, a principal component analysis (PCA) of the infectome revealed a clear distinction between infected vs. non-infected macrophages (component 1, 53.4%), as well as temporal and strain-specific responses (component 2, 4.2%) (Fig. 1H).

### Host response to *S*. Typhimurium infection reveals temporal and strain-specific regulation

During infection, dynamics proteome changes are measured from both the host and pathogen perspectives to define how the host cells protect themself from invasion and how the pathogen evades or adapts to the host response. Here, we focus on infection from the host’s perspective by profiling changes in protein abundance over time and in the presence of the different *S*. Typhimurium strains. By PCA, we observed three distinct clusters of samples based on time and strain (Fig. 2A). Specifically, the early time points of infection (0 and 0.5 hpi) for non-infected, WT-infected, and Δ*spi1*-infected samples clustered. Whereas the early (0 and 0.5 hpi) and mid (1 and 2 hpi) time points cluster for *spi2-*defective -infected samples, along with the mid time points (1 and 2 hpi) for WT-infected and Δ*spi1*-infected samples, and mid (1 and 2 hpi) and late (6 hpi) time points for the non-infected samples. Finally, we observed a clear separation for the late time point (6 hpi) for all infected samples, indicating at 6 hpi, we observe a consistent host response in the presence of the bacterial strains, which can be teased apart earlier on in the infection process, depending on the strain. These data support the distinct roles of SPI-1 and SPI-2 in bacterial invasion and replication, respectively, within macrophages, and clearly define a time-dependent host response.

**Fig 2.**
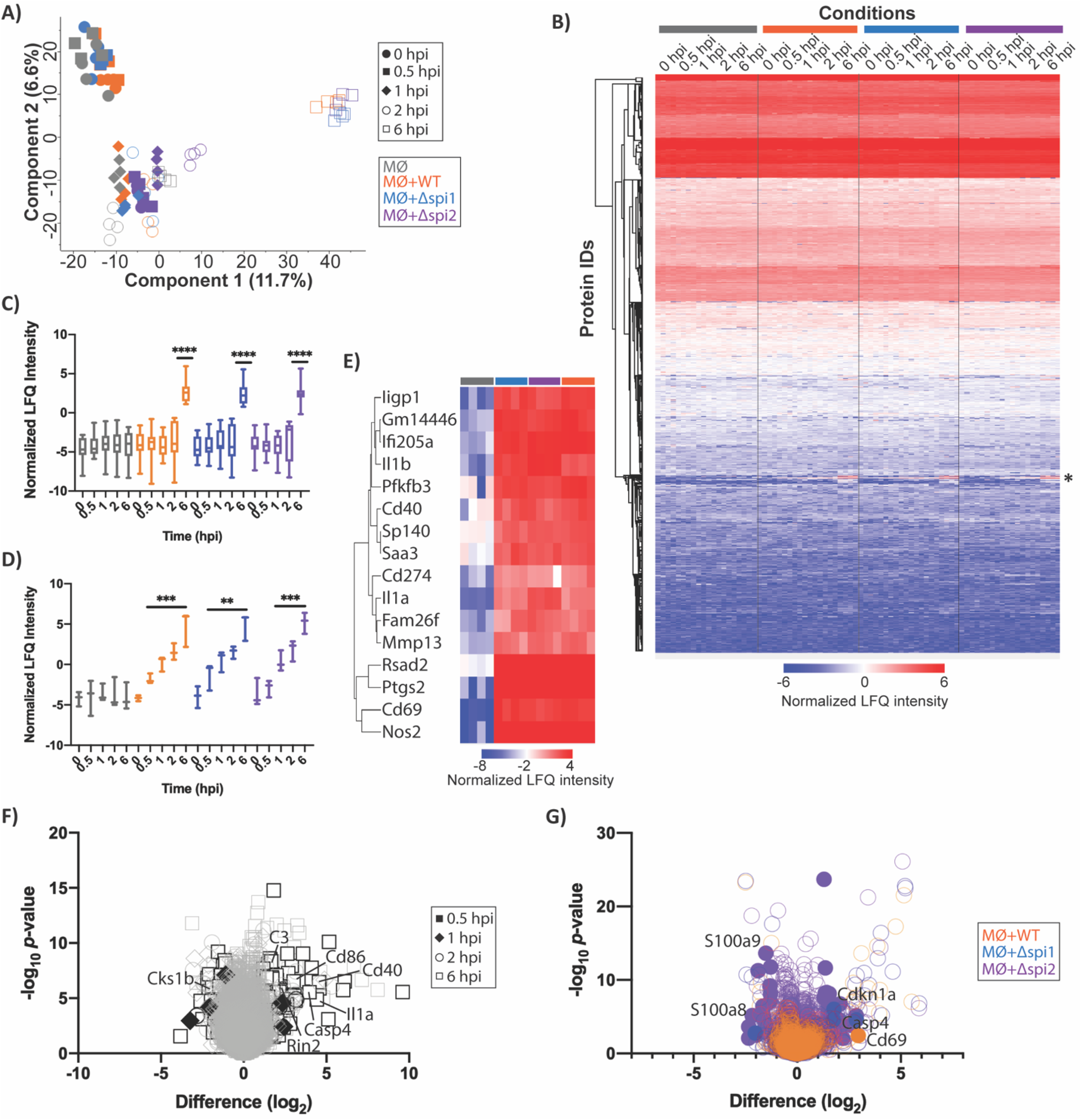
Host-specific response to *S*. Typhimurium infection. (A) Principal component analysis of host proteins over time course of infection (0, 0.5, 1, 2, 6 hpi) with different bacterial strains (WT, Δ*spi1, spi2-*defective). (B) Hierarchical clustering by Euclidean distance for protein IDs based on normalized LFQ intensity. *Represents time-dependent profile observed among infected host samples. (C) Tracing of time-dependent host response (6 hpi) to *S*. Typhimurium infection (n = 13 proteins). (D) Tracing of increasing time-dependent host response (0, 0.5, 1, 2, 6 hpi) to *S*. Typhimurium infection (n = 3 proteins). For box plots, statistical analysis by Student’s *t*-test: \*\**p*- value < 0.005; \*\*\**p*-value < 0.0005; \*\*\*\**p*-value < 0.00001. (E) Hierarchical clustering by Euclidean distance for immune-associated proteins of interest (n = 16 proteins). (F) Stacked volcano plot for time-dependent host response. Significantly different proteins unique to each time point are outlined/highlighted over commonly identified proteins (grey). (G) Stacked volcano plot for bacterial strain-dependent host response. Significantly different proteins unique to each bacterial strain are highlighted (closed circles) over commonly identified proteins (open circles). For volcano plots, statistical analysis by Student’s *t*-test (*p*-value ≤ 0.05); FDR = 0.05; S0=1.

Next, clustering by protein IDs via hierarchical clustering highlighted protein groups associated with the temporal host response (Fig. 2B). A closer look at host proteins with increased abundance over the time course of infection revealed a subset of proteins with significant increases in abundance at 6 hpi regardless of the bacterial strain applied to the macrophage (Fig. 2C). These proteins included induction of host immunity (e.g., IL-1⍺, Nos2, Cd40) and defense response (e.g., Sp140) (Supp. Table 2). Next, we noticed a trend for proteins to significantly increase in abundance over the time course of infection, including innate immune effector (e.g., Il1B) (Fig. 2D; Supp. Table 3) and immune response-associated proteins (i.e., Sp140, Pfkfb3). Hierarchical clustering of immune-associated proteins demonstrated clustering by Euclidean distance (Fig. 2E).

To explore significant changes in abundance of proteins unique to each of the collected time points, we generated stacked volcano plots to highlight time-specific host responses (Fig. 2F). This analysis identified significantly different proteins unique to each time point, including 1 hpi (five proteins), 2 hpi (four proteins), and 6 hpi (153 proteins) (Supp. Table 4). Notably, at 0.5 hpi, no significant changes in abundance were observed. As anticipated, at 6 hpi, we observed the largest change in unique host protein response with involvement of many immune associated proteins (e.g., Cd86, Cd40, Casp4, C3, IL-1⍺), as well as proteins involved in signaling cascades (e.g., Cks1b) and regulatory networks (e.g., Rin2), demonstrating a specific and complex response to bacterial infection over time. Taken together, these data demonstrate time-specific responses induced within the host during infection, regardless of the bacterial strain, and represents key points for further exploration as potential inhibitory or stimulatory host targets in defense against invading bacterial pathogens.

Next, to assess unique host responses dependent on the *S*. Typhimurium strain, we generated stacked volcano plots to highlight strain-specific significantly different proteins (Fig. 2G). For each strain, we observed unique host cell expression pattern (Supp. Table 5). For example, in the presence of WT, macrophages uniquely produced Cd69, an early activation antigen, suggesting activation of this host protein is related to T3SS function. For Δ*spi1* infection, we observed four uniquely produced proteins with diverse roles in cellular processes, including a signaling inhibitor (Cdkn1a). Notably, given the distinction of WT to invade macrophages vs. entry only by phagocytosis for Δ*spi1* due to a lack of invasion properties, these protein differences may be linked to unique intracellular locations of the bacteria within the macrophage. For *spi2-*defective infection, we observed the unique production of 49 proteins (28 with increased abundance; 21 with decreased abundance), indicating specific host defense programs to *S*. Typhimurium in the absence of bacterial cell replication, including specific reduction in pro-inflammatory signals (e.g., S100a9, S100a8) and induction of inflammasome components (e.g., Casp4). Overall, these bacterial strain-specific responses demonstrate remodeling of the host proteome in response to the presence or absence of pathogenicity islands and provide new insight into the complementary roles SPI-1 and SPI-2 play in *Salmonella* infection.

### Co-regulation of novel infection-associated proteins and *Salmonella* pathogenicity islands

Here, we aimed to identify novel infection-associated proteins influenced by the presence or absence of the pathogenicity islands during a time course of infection of primary macrophages. To identify novel infection-associated proteins, we compared the *in vitro* cellular proteomes and secretomes of Δ*spi1* and *spi2-*defective to the respective infectomes, relative to WT. This approach defines proteins with increased abundance during infection and identifies a subset of previously uncharacterized proteins with posssible roles in virulence. We selected candidates for functional follow-up based on abundance changes, predicted secretion or membrane location for interaction with the host cell, potential roles in virulence, and regulation influenced by SPI-1 and/or SPI-2. As expected, we also identified known virulence factors, serving as positive controls for our approach, and allowed for further characterization of their roles in *Salmonella* virulence.

A comparison of WT-Δ*spi1* cellular proteome vs. infectome identified 28 proteins with higher abundance during infection in the absence of *spi1* (Fig. 3A). Of these proteins, 17 are associated with SPI-1 and SPI-2. We prioritized two proteins (SL1344_1263, SL1344_3112) which are produced *in vitro,* displaying higher abundance during infection, and potentially interact with the host via localization (i.e., membrane) and secretion (i.e., signal peptide) (Supp. Table 6).

**Fig 3.**
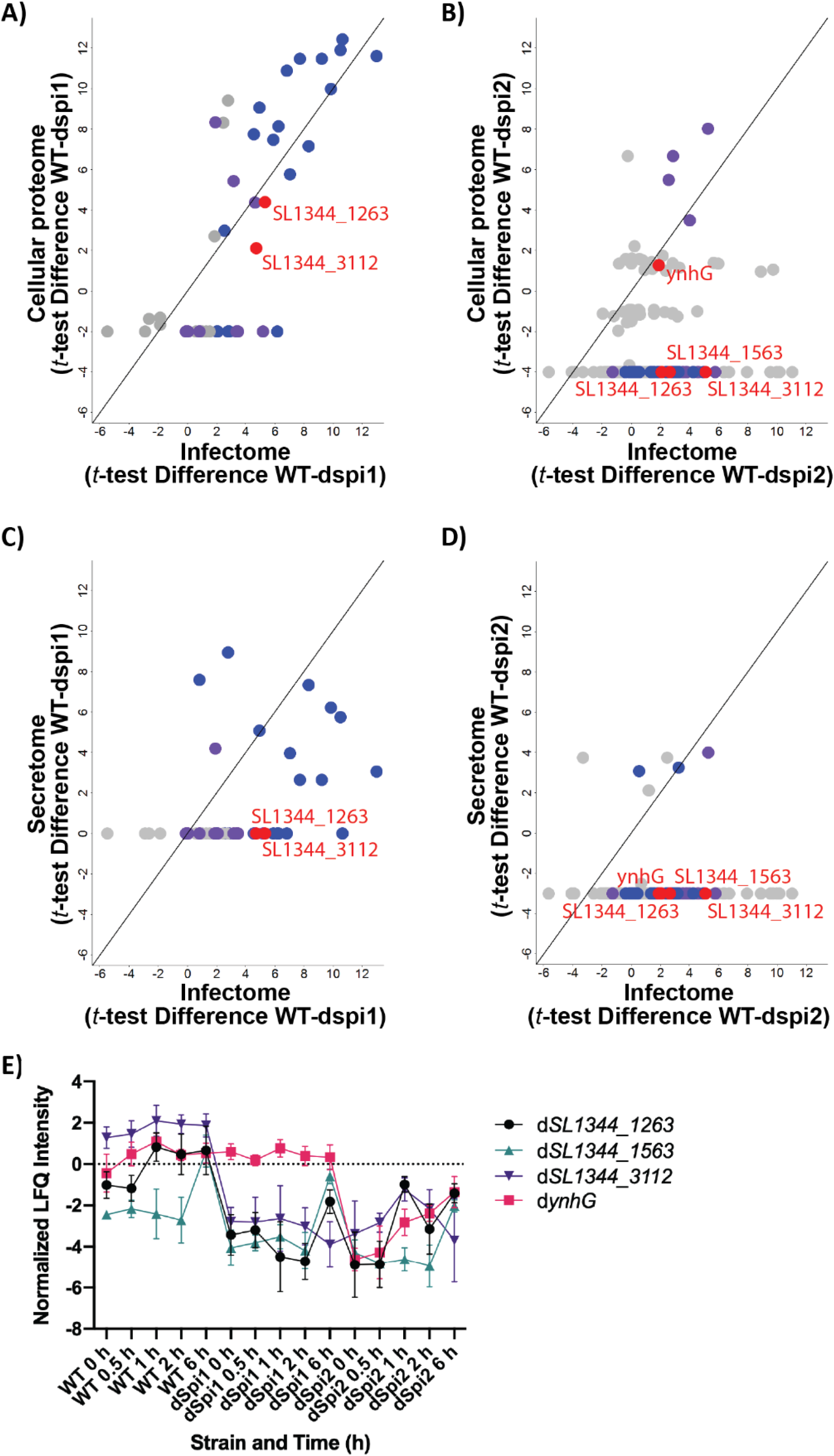
Identification of novel infection-associated bacterial proteins regulated by the presence of *spi1* or *spi2*. (A) Comparison of significantly different bacterial proteins identified from cellular proteome and infectome profiling between WT and Δ*spi1*. (B) Comparison of significantly different bacterial proteins identified from cellular proteome and infectome profiling between WT and *spi2-*defective. (C) Comparison of significantly different bacterial proteins identified from secretome and infectome profiling between WT and Δ*spi1*. (D) Comparison of significantly different bacterial proteins identified from secretome and infectome profiling between WT and *spi2-*defective. For plots A-D, line represents no change in protein abundance between *in vitro* (cellular proteome or secretome) vs. infectome profiling. Spi1-encoded and -effector proteins (blue); Spi2-encoded and -effector proteins (purple); candidate novel infection-associated bacterial proteins (red). Statistical analysis by Student’s *t*-test (*p*-value ≤ 0.05); FDR = 0.05; S0=1. (E) Tracing of time- and strain-dependent protein abundance of candidate infection-associated bacterial proteins.

Of the remaining nine proteins, roles in motility, biosynthesis, and metabolism were described based on Gene Ontology Biological Process (GOBP). A comparison of WT*-spi2-*defective cellular proteome vs. infectome identified 316 proteins with higher abundance during infection in the disruption of *spi2,* supporting a larger discrepancy between the impact of infection from WT vs. SPI-2, compared to WT vs. SPI-1 (Fig. 3B). Of these proteins, 32 are associated with SPI-1 and SPI-2, two proteins were also detected in the SPI-1 comparisons (SL1344_1263, SL1344_3112) and two were hypothetical proteins with signal peptides (YnhG, SL1344_1563) (Supp. Table 7). Of the remaining 280 proteins, roles in biosynthesis, metabolism, motility, transcription, translation, and enzymatic activity are described by GOBP, suggesting opportunity for curated future exploration. Secretome analyses supported these findings (Fig. 3C & 3D; Supp. Table 8 & 9).

Time-resolved SPI-1 and SPI-2 dependent protein profiles revealed abundance changes for SL1344_1563 (i.e., increased abundance at 6 hpi), strain- and time-associated response for SL1344_1263 (i.e., influenced by absence of SPI-1 and SPI-2 over the time course of infection), as well as strain-specific responses (i.e., SL1344_3112 regulated by the absence of SPI-1 and SPI- 2, and *ynhG* with a delayed response when SPI-2 is absent) (Fig. 3E). Profiling of novel infection-associated bacterial protein abundance in the cellular proteome and secretome, align with these infectome observations in WT, Δ*spi1*, and *spi2-*defective strains, supporting potential co-regulatory roles (Supp. Fig. 1). Taken together, our comparison of proteins produced during *in vitro* bacterial growth vs. co-infection with primary macrophages revealed a sub-set of novel infection-associated proteins with regulatory patterns influenced over the time course of infection and dependent on the pathogenicity islands in *Salmonella*.

### Characterization of novel infection-associated bacterial proteins reveals roles in intracellular lifestyle and virulence

To evaluate whether the novel infection-associated bacterial proteins impact host cell response, we generated bacterial deletion mutants and performed *in vitro* and *in vivo* loss-of-function infection experiments. Our initial analysis monitored induced cell death by lactate dehydrogenase (LDH) release over the time course of infection for the *Salmonella* strains. We report similar macrophage cytotoxicity for Δ*spi1* and uninfected cells, as anticipated given the inability of Δ*spi1* to invade, along with similar trends in macrophage cytotoxicity amongst WT, *spi2*-defective, and novel infection-associated proteins (Fig. 4A). Next, we performed an association, invasion, and replication (AIR) assay to monitor difference in functions among the tested strains with Δ*spi1* and *spi2-*defective serving as controls for invasion and replication, respectively (Fig. 4B). We observed similar rates of association among the strains compared to WT with significant increase in association for Δ*spi1*. For invasion, we observed a significant reduction in Δ*spi1*, ΔSL1344_3112, and ΔSL1344_1263, supporting connections with SPI-1 regulation, as confirmed by infectome profiling presented above, as well as a reduction in invasion for *spi2-*defective. Lastly, we showed a significant reduction in replication for *spi2-*defective, as anticipated, and ΔSL1344_1563, further supporting our observation of SL1344_1563 and co- regulation with SPI-2.

**Fig 4.**
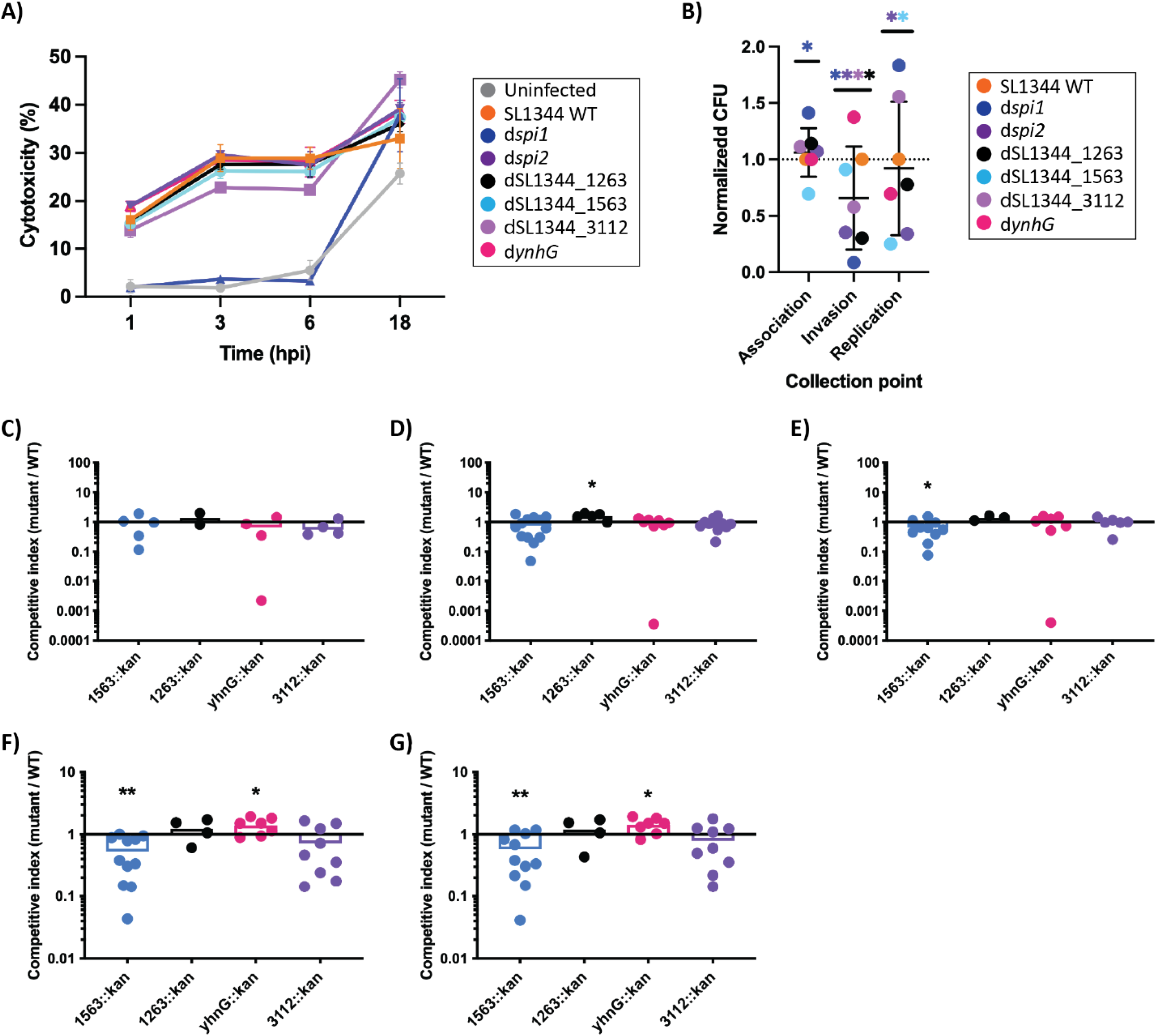
*In vitro* intracellular survival assays and *in vivo* virulence assays. (A) Cytotoxicity assay by LDH of macrophage infected with *S*. Typhimurium strains (WT; Δ*spi1; spi2-*defective; ΔSL1344_1263; ΔSL1344_1563; ΔSL1344_3112; Δ*ynhG*) over a time-course of infection (0, 1, 3, 6, 18 hpi). (B) Association, replication, and invasion assay of S. Typhimurium strains (WT; Δ*spi1; spi2-*defective; ΔSL1344_1263; ΔSL1344_1563; ΔSL1344_3112; Δ*ynhG*). (C) Murine model of infection – Ileum: competitive index assay. (D) Murine model of infection – Cecum: competitive index assay. (E) Murine model of infection – Colon: competitive index assay. (F) Murine model of infection – Spleen: competitive index assay. (G) Murine model of infection – Liver: competitive index assay. Competitive index assays performed with six-week-old female C57BL/6 mice in groups of 10 following oral gavage with WT and mutant 1:1 in the typhoid model. Experiment performed in biological triplicate; \**p*-value <0.05; \*\**p*-value <0.01. LOD = limit of detection.

To examine the contribution of the candidate proteins in virulence, we used a typhoid model of murine infection by competitive index with WT, causing moderate colonization of the gut and systemic dissemination [49]. In tissues representing initial infection (i.e., ileum, cecum, colon), commonly influenced by bacterial strain invasion, we observed no significant change in abundance for any of the strains in the ileum (Fig. 4C) but a significant increase in abundance of the ΔSL1344_1263 strain, relative to WT, in the cecum, further supporting a role of co-regulation between SL1344_1263 and SPI-1 (Fig. 4D). We also observed a significant decrease in abundance of ΔSL1344_1563, compared to WT, in the colon (Fig. 4E). Next, we assessed bacterial load in tissues associated with systemic infection (i.e., spleen, liver) and survival fitness of the bacterium (i.e., replication). Similar to our observation in the colon, we showed a significant decrease in bacterial load counts of ΔSL1344_1563 compared to WT in the spleen (Fig. 4F) and liver (Fig. 4G), as well as a significant increase in abundance of Δ*ynhG* in the spleen (Fig. 4F) and liver (Fig. 4G), supporting roles in bacterial replication and co-regulation with SPI-2. Notably, ΔSL1344_3112 did not show any significant differences in virulence within the tested models and was not used for further analyses. Importantly, of the bacterial strains evaluated, only ΔSL1344_1563 demonstrated a reduction in virulence, whereas the other strains showed heightened virulence in the absence of the proteins, suggesting co-regulation with SPI-1 and SPI- 2 in positive and negative manners. Overall, these data reveal novel T3SS effector roles for infection-associated proteins in bacterial survival and virulence in the host.

### Time-dependent bacterial protein production confirms proteome profiling

Based on our *in vitro* and *in vivo* data, we assessed the production of the novel T3SS effector proteins, proteins *in vitro* (i.e., during bacterial growth). Here, FLAG-tagged effector proteins under regulation of the native promoter allowed for tracking and quantifying production over a time course of bacterial growth (Fig. 5A). We observed a time-dependent increase in protein production for SL1344_1563 over 6 h of bacterial growth, corroborating with the time-dependent infectome protein profile (Fig. 3) and roles in virulence during systemic infection (Fig. 4). For SL1344_1263, we observed an early induction of the protein at 1 h, which decreased over time, aligning with infectome abundance (Fig. 3) and co-regulation by SPI-1, and early colonization of the gut (Fig. 4). Lastly, we profiled YnhG production over time and observed consistent abundance throughout the growth curve, suggesting on/off production influenced by the presence of the host. We confirmed protein production by quantifying the proportion of cells with FLAG production (Fig. 5B) and Western blot (Fig. 5C) and showed increased secretion of SL1344_1563 in culture supernatant over time (Fig. 5D). Together, profiling of FLAG-tagged protein abundance within *Salmonella* validates infectome and virulence characterization and supports SPI-1 and SPI-2 patterns of co-regulation and provides evidence of protein secretion.

**Fig 5.**
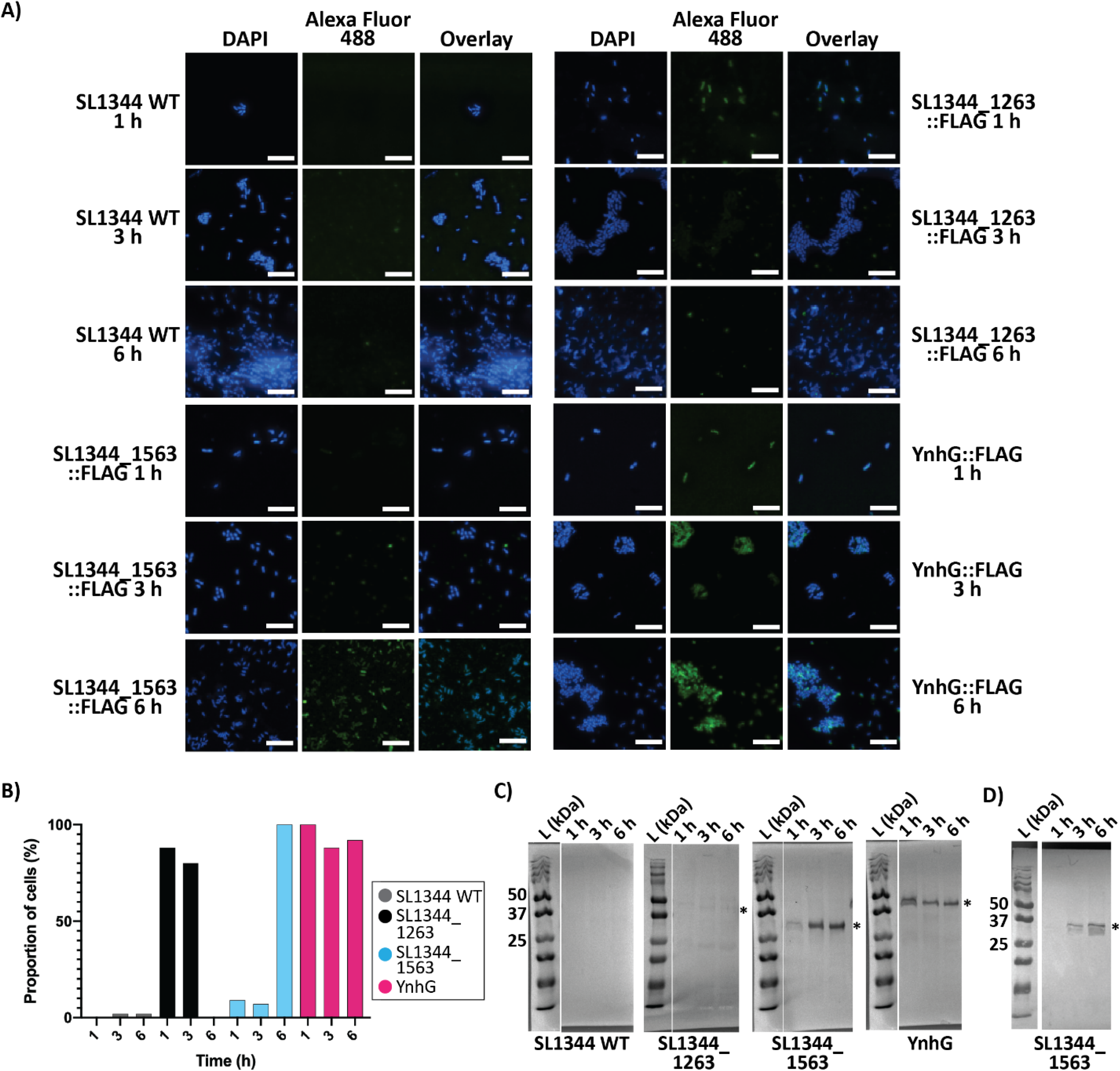
Characterization of novel infection-associated bacterial proteins. (A) Fluorescence microscopy of FLAG-tagged *S*. Typhimurium strains (SL1344_1263; SL1344_1563; YnhG). Images captured at 1, 3, and 6 h with DAPI and Alexa Fluor 488 (FLAG). Scale bar 10µM. (B) Bar plot of percent of detected bacterial cells expressing FLAG assessed from fluorescence micrographs. (C) Western blot of cell pellet collected at 1, 3, and 6 h with anti-FLAG antibody. Protein sizes: SL1344_1263 = 34.8 kDa, SL1344_1563 = 28.4 kDa, and YnhG = 36.1 kDa. (D) Western blot of supernatant collected at 1, 3, and 6 h with anti-FLAG antibody. Protein sizes: SL1344_1563 = 28.4 kDa. *denotes expected protein size. For plots A-D, WT strain does not contain FLAG-tagged protein, used as a negative control. L = Ladder; 50, 37, and 25 kDa.

### Bacterial protein secretion and interactions within host cells as molecular checkpoints for perturbation of infection-associated proteins

We next investigated production and secretion of the infection-associated proteins in the presence of the host. We showed that both SL1344_1563 and YnhG are produced in the presence of macrophages at 6 hpi (Fig. 6A). Whereas SL1344_1563 emerged as a bacterium affiliated within the macrophage, supporting its role as a transmembrane protein involved in amino acid transport [50], YnhG was secreted outside the macrophage. Production of SL1344_1263 was not detected.

**Fig 6.**
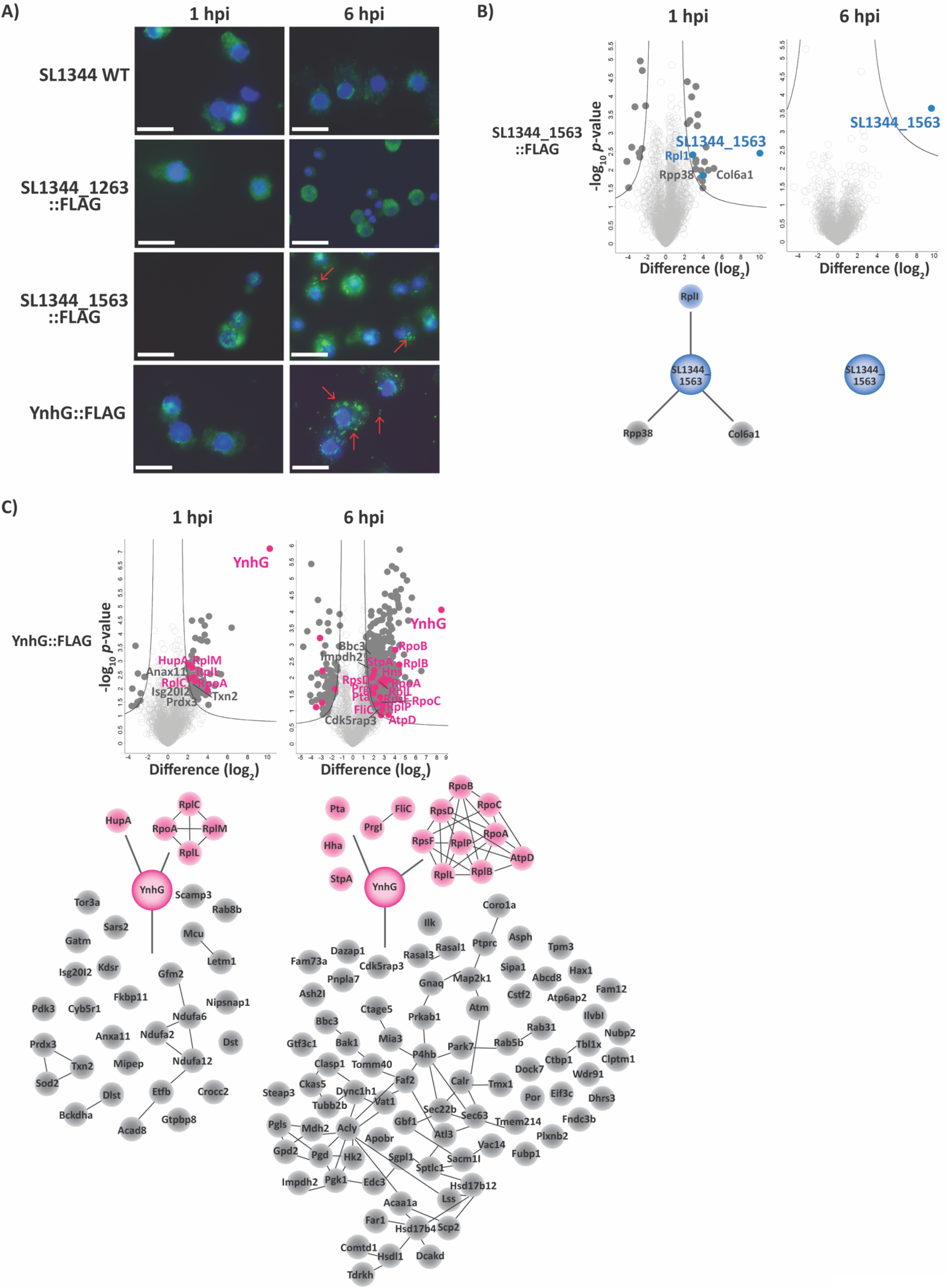
Interactome of novel infection-associated bacterial proteins. (A) Fluorescence microscopy of FLAG-tagged *S*. Typhimurium strains (SL1344_1263; SL1344_1563; YnhG) co-cultured with macrophage (MOI 100:1). Images captured at 1 and 6 h with DAPI and Alexa Fluor 488 (FLAG). Scale bar 10µM. (B) Interactome volcano plots of SL1344_1563::FLAG at 1 and 6 hpi with macrophage (MOI 100:1) (top) and interactome mapping of bacterial (blue) and host (grey) proteins. (C) Interactome volcano plots of YnhG::FLAG at 1 and 6 hpi with macrophage (MOI 100:1) (top) and interactome mapping of bacterial (pink) and host (grey) proteins. For volcano plots, statistical analysis by Student’s *t*-test (*p*-value ≤ 0.05); FDR = 0.05; S0=1. Lines connecting bacterial and host nodes within interactome maps referenced from STRING database interaction mapping. Notably, autofluorescence of macrophages was observed and minimized with media selection; positive secretion patterns were determined by visualization of bacterial cells (i.e., SL1344_1563 and YnhG).

To detect interactions between novel *Salmonella* effector and host proteins, we performed an immunoprecipitation assay during macrophage infection using bacterial strains expressing FLAG-tagged effectors. Interactome profiling of SL1344_1563 revealed few unique interactions with the host, including ribonuclease P (Rpp38) and collagen alpha-1 chain (Col6a1) at 1 hpi (Fig. 6B; Supp. Table 10). We also found that SL1344_1563 interacted with a bacterial 50S ribosomal protein (RplI) at 1 hpi and no interactions were observed at 6 hpi. For YnhG, the interaction was more diverse and widespread. For example, at 1 hpi, YnhG interacted with five bacterial proteins, including ribosomal proteins (RplC, RplM, RplL), RNA polymerase (RpoA), and a DNA-binding protein (HupA), and 30 host proteins (Fig. 6C; Supp. Table 11). Network analysis of the host proteins using the STRING database revealed known and predicted connections among the proteins, highlighting targeted immune functions (e.g., Anxa11, Isg20I2) and defense response (e.g., Txn2, Prdx3). Similarly at 6 hpi, YnhG interacted with 15 bacterial proteins with roles in translation and transcription (i.e., RpoB, RpsD, RpoC, RpoA, RpsF, RplP, RplL, RplB, AtpD), motility (i.e., FliC), biosynthesis (i.e., Pta), and pathogenesis (i.e., PrgI, Hns, StpA). YnhG also interacted with 85 host proteins, including proteins of the immune response (i.e., Impdh2) and numerous programmed cell death (i.e., Bbc3, Cdk5rap3), suggesting a role for YnhG in regulating the host’s ability to combat infection through programmed cell death (Supp. Table 12). Further investigation to tease apart direct and indirect interactors, as well as perturbation of these interactions between the host and pathogen present potential opportunities for novel anti-virulence strategies to combat *Salmonella* infection.

## Discussion

In this study, we profile differences of the bacterial proteome and infectome in consideration of both SPI-1 and SPI-2 over a time course of infection in primary macrophages. We define time-dependent and strain-specific responses of the host to *Salmonella* infection and uncover unique patterns of cellular regulation dependent upon these parameters. In addition, we report novel infection-associated bacterial proteins by comparing abundance during bacterial growth *in vitro* (i.e., cell culture) to co-culture with primary macrophages and we demonstrate co-regulation with SPI-1 and/or SPI-2 for these effectors. We dissect the role of the infection-associated proteins in bacterial virulence using *in vitro* (i.e., LDH & AIR assays) and *in vivo* (i.e., murine infection) models by generation of bacterial knockout strains, and we observe temporal production differences associated with induction of SPI-1 and SPI-2 *in vitro*. Moreover, we characterize the role of two infection-associated proteins, SL1344_1563 and YnhG, within the macrophages by visualizing their production and secretion, as well as generating a network of novel interactions between the pathogen and host proteins. Overall, our approach identifies and characterizes novel T3SS effectors regulated by SPI-1 and SPI-2. The method is universal with potential in diverse biological systems to discover new infection-associated proteins and delinate their roles in cellular regulation, as well as interactions with the host that are critical for infection. Such connectivity between the host and pathogen suggests opportunity for perturbation to disrupt the observed interactions as a method of regulating bacterial virulence and controlling progression of disease.

### New insights for a known virulence-associated bacterial protein, SL1344_1563

Previous structural and biochemical characterization of the periplasmic-binding protein component of a D-alanine ABC transporter, deemed DalS (also known as SL1344_1563 in strain SL1344), was co-regulated with the SPI-2 virulence locus through direct activation of response regulator, SsrB [50]. In the current study, we observe similar co-regulation with SPI-2, along with abundance of immune related proteins during infection of primary macrophages. We observe a fitness defect for Δ*dalS in vivo* for overall survival and significant differences in competitive index in the liver and cecum [50]. We build upon these observations by reporting significant reduction in competitiveness of the ΔSL1344_1563 strain in the spleen and liver, supporting a role in systemic infection, which aligns with the ability of bacteria to survive and replicate in the host, as associated with the SPI-2 virulence locus. An additional study explored the protective power of DalS for *Salmonella* from D-amino acid oxidase-dependent killing in neutrophils [54]. Here, we provide additional characterization of links between DalS and the host immune system by defining a role in macrophage infection at the protein level, generating protein expression profiles within the bacteria as well as during co-culture with macrophages, and determining unique interactions between DalS and the host. Specifically, we report the interaction between SL1344_1563 and the bacterial protein, RplI, a 50 S ribosomal protein associated with translation, as well as two host proteins, including Rpp38, a ribonuclease P protein involved in rRNA processing, and Col6a1, a collagen alpha-1(VI) chain protein involved in cell adhesion and cellular response to amino acid stimulus [55]. Col6a1 was previously linked to an organ atlas of systemic bacterial infection with group A *Streptococcus*, as well as a recent connection of host infection with SARS-CoV-2 as a druggable target, implicating important roles in disease [56, 57]. Further experimentation into the importance of the SL1344_1563-Col6a1 interaction is underway as a potential target to interrupt the interaction between pathogen and host during infection. For example, emphasis on the benefits of anti-virulence strategies to combat microbial infections have taken hold based on the potential to reduce virulence factor effectiveness without directly killing the pathogen, providing a selective therapeutic advantage without affecting commensal bacteria (e.g., microbiota) [58]. Such approaches lead to enhanced killing by the host’s immune system and an overall reduction in selective pressure towards the development of antimicrobial resistance. Moreover, anti-virulence strategies may require rational drug design to specifically target the interaction between the host and pathogen proteins, a process fully supported through developments in mass spectrometry-based proteomics [59].

### New insights for ***YnhG –*** an exported bacterial protein with implications in intercellular signaling and clearance of infection

YnhG is classified as a hypothetical exported protein involved in the peptidoglycan biosynthetic process as defined by GOBP. The presence of a signal peptide and LysM domain, a highly-conserved carbohydrate binding module recognizing polysaccharides containing N-acetylglucosamine residues found in the bacterial cell wall, support its roles in building the bacterial cell surface, a critical point of interaction with the host [60]. Molecules exposed on bacterial surfaces elicit innate immune response through pattern recognition receptors, induction of cytokines and apoptosis, as well as activation of antimicrobial activity [61]. In addition, Lys-M domain-containing proteins of the host (e.g., LysMD1, LysMD2, LysMD3, and LysMD4) are assumed to be involved in immune responses against bacterial infection [62, 63]. We also observed interaction of YnhG with several bacterial proteins at early (1 hpi) and late (6 hpi) time points of co-infection with connections to pathogenesis, including StpA and PrgI, suggesting a role of this protein in *Salmonella* virulence. In addition, YnhG interacts, either directly or indirectly with a plethora of host proteins, suggesting additional intercellular interaction with the host through secretion from the bacterial cell. Deletion of *ynhG* resulted in increased bacterial load counts in the spleen and liver following competitive index assays with a slight reduction in colonization of the ileum. These findings suggest a role for YnhG negative regulation of the host’s immune response against the pathogen and therefore, strategies to enhance production of YnhG may alter disease outcome.

## Conclusion

Overall, we report time-dependent and strain-specific proteomic responses of the host to *Salmonella* infection, and we discover and characterize new T3SS effector proteins regulated by SPI-1 and SPI-2 with implications in drug design. We provide a new strategy for defining the intricate relationship between a host and pathogen from a global and unbiased perspective to advance our understanding of infectious diseases and reveal new opportunities to combat infection.

## Ethics Statement

For murine models of infection, experiments were performed in accordance with the guidelines of the Canadian Council on Animal Care and the University of British Columbia (UBC) Animal Care Committee (certificate A17-0228).

## Availability of Data

The mass spectrometry proteomics data have been deposited in the PRIDE partner repository for the ProteomeXchange Consortium with the data set identifier: PXD027653

Reviewer login

Password: CegwyaGp

## Acknowledgements

The authors thank members of the Geddes-McAlister, Meissner, Finlay, and Khursigara labs for critical reading of the manuscript and insightful comments. We thank Dr. Jonathan Krieger (Bioinformatics Solutions Inc.) for operating the mass spectrometer for interactome experiments. The authors also thank Drs. David Holden, Sophie Helaine, and Jay Hinton for their helpful insights and suggestions.

## Contributions

J.G.-M. & F.M. perceived the project. J.G.-M., A.S., S.L.V., E.J.R., C.M.K., B.R., B.B.F., & F.M. planned experiments. J.G.-M., A.S., S.L.V., J.L.R., S.E.W., L.G., & E.J.R. performed experiments. J.G.-M., A.S., S.L.V., E.J.R., C.M.K., & F.M. analyzed and interpreted data. J.G.- M., S.L.V., B.M., & E.J.R. generated figures. J.G.-M. wrote the manuscript. All authors read, edited, and approved the submitted manuscript.

## Supplementary files

Supplementary Figure 1: Co-regulation of candidate infection-associated proteins *in vitro*.

**Supplementary Table 1: Primers used in this study.**

**Supplementary Table 2: Time-dependent host proteins (6 hpi).**

**Supplementary Table 3: Progressive time-dependent host proteins.**

**Supplementary Table 4: Unique time-dependent significantly different host proteins.**

**Supplementary Table 5: Unique strain-specific significantly different host proteins.**

**Supplementary Table 6: Candidate infection-associated bacterial proteins from infectome vs. cellular proteome comparison of dspi-1.**

**Supplementary Table 7: Candidate infection-associated bacterial proteins from infectome vs. cellular proteome comparison of dspi-2.**

**Supplementary Table 8: Candidate infection-associated bacterial proteins from infectome vs. secretome comparison of dspi-1.**

**Supplementary Table 9: Candidate infection-associated bacterial proteins from infectome vs. secretome comparison of dspi-2.**

**Supplementary Table 10: Interactome of SL1344_1563::FLAG-tag at 1 hpi. Supplementary Table 11: Interactome of YnhG::FLAG-tag at 1 hpi.**

**Supplementary Table 12**: **Interactome of YnhG::FLAG-tag at 6 hpi**

## References

1. Sánchez-Vargas FM, Abu-El-Haija MA, Gómez-Duarte OG. *Salmonella* infections: An update on epidemiology, management, and prevention. Travel Medicine and Infectious Disease. 2011. doi:10.1016/j.tmaid.2011.11.001

2. Darwin KH, Miller VL. Molecular basis of the interaction of *Salmonella* with the intestinal mucosa. Clin Microbiol Rev. 1999. doi:10.5665/sleep.2450

3. Birmingham CL, Smith AC, Bakowski MA, Yoshimori T, Brumell JH. Autophagy controls *Salmonella* infection in response to damage to the *Salmonella*-containing vacuole. J Biol Chem. 2006. doi:10.1074/jbc.M509157200

4. Wood MW, Jones MA, Watson PR, Siber AM, McCormick BA, Hedges S, et al. The secreted effector protein of *Salmonella* dublin, SopA, is translocated into eukaryotic cells and influences the induction of enteritis. Cell Microbiol. 2000. doi:10.1046/j.1462-5822.2000.00054.x

5. Patel JC, Galán JE. Manipulation of the host actin cytoskeleton by *Salmonella* - All in the name of entry. Current Opinion in Microbiology. 2005. doi:10.1016/j.mib.2004.09.001

6. McGhie EJ, Brawn LC, Hume PJ, Humphreys D, Koronakis V. *Salmonella* takes control: effector-driven manipulation of the host. Current Opinion in Microbiology. 2009. doi:10.1016/j.mib.2008.12.001

7. Buckner MMC, Croxen M, Arena ET, Finlay BB. A comprehensive study of the contribution of *Salmonella enterica* serovar Typhimurium SPI2 effectors to bacterial colonization, survival, and replication in typhoid fever, macrophage, and epithelial cell infection models. Virulence. 2011. doi:10.4161/viru.2.3.15894

8. Brumell JH, Tang P, Zaharik ML, Finlay BB. Disruption of the *Salmonella*-containing vacuole leads to increased replication of *Salmonella enterica* serovar Typhimurium in the cytosol of epithelial cells. Infect Immun. 2002. doi:10.1128/IAI.70.6.3264-3270.2002

9. Haraga A, Ohlson MB, Miller SI. *Salmonellae* interplay with host cells. Nature Reviews Microbiology. 2008. doi:10.1038/nrmicro1788

10. Di Marzio M, Shariat N, Kariyawasam S, Barrangou R, Dudley EG. Antibiotic resistance in *Salmonella enterica* serovar Typhimurium associates with CRISPR sequence type. Antimicrob Agents Chemother. 2013. doi:10.1128/AAC.00913-13

11. Bruno VM, Hannemann S, Lara-Tejero M, Flavell RA, Kleinstein SH, Galán JE. *Salmonella* Typhimurium type III secretion effectors stimulate innate immune responses in cultured epithelial cells. PLoS Pathog. 2009. doi:10.1371/journal.ppat.1000538

12. Jennings E, Thurston TLM, Holden DW. *Salmonella* SPI-2 Type III Secretion System Effectors: Molecular Mechanisms And Physiological Consequences. Cell Host and Microbe. 2017. doi:10.1016/j.chom.2017.07.009

13. Van Der Heijden J, Finlay BB. Type III effector-mediated processes in *Salmonella* infection. Future Microbiology. 2012. doi:10.2217/fmb.12.49

14. Kröger C, Colgan A, Srikumar S, Händler K, Sivasankaran SK, Hammarlöf DL, et al. An infection-relevant transcriptomic compendium for *Salmonella enterica* serovar Typhimurium. Cell Host Microbe. 2013;14: 683–695. doi:10.1016/j.chom.2013.11.010

15. Figueira R, Watson KG, Holden DW, Helaine S. Identification of *Salmonella* pathogenicity island-2 type III secretion system effectors involved in intramacrophage replication of *S. enterica* serovar Typhimurium: Implications for rational vaccine design. MBio. 2013. doi:10.1128/mBio.00065-13

16. Jiang L, Wang P, Song X, Zhang H, Ma S, Wang J, et al. *Salmonella* Typhimurium reprograms macrophage metabolism via T3SS effector SopE2 to promote intracellular replication and virulence. Nat Commun. 2021;12: 1–18. doi:10.1038/s41467-021-21186-4

17. Métris A, Sudhakar P, Fazekas D, Demeter A, Ari E, Olbei M, et al. SalmoNet, an integrated network of ten *Salmonella enterica* strains reveals common and distinct pathways to host adaptation. npj Syst Biol Appl. 2017;3: 1–10. doi:10.1038/s41540-017-0034-z

18. Aebersold R, Mann M. Mass-spectrometric exploration of proteome structure and function. Nature. 2016;537: 347–355. doi:10.1038/nature19949

19. Sukumaran A, Coish JM, Yeung J, Muselius B, Gadjeva M, MacNeil AJ, et al. Decoding communication patterns of the innate immune system by quantitative proteomics. Journal of Leukocyte Biology. 2019. doi:10.1002/JLB.2RI0919-302R

20. Sukumaran A, Woroszchuk E, Ross T, Geddes-McAlister J. Proteomics of host-bacterial interactions: new insights from dual perspectives. Can J Microbiol. 2020; 1–43.

21. Liu Y, Zhang Q, Hu M, Yu K, Fu J, Zhou F, et al. Proteomic analyses of intracellular *Salmonella enterica* serovar Typhimurium reveal extensive bacterial adaptations to infected host epithelial cells. Infect Immun. 2015;83: 2897–2906. doi:10.1128/IAI.02882-14

22. Liu Y, Yu K, Zhou F, Ding T, Yang Y, Hu M, et al. Quantitative Proteomics Charts the Landscape of *Salmonella* Carbon Metabolism within Host Epithelial Cells. J Proteome Res. 2017;16: 788–797. doi:10.1021/acs.jproteome.6b00793

23. Rogers LD, Brown NF, Fang Y, Pelech S, Foster LJ. Phosphoproteomic analysis of *Salmonella*-infected cells identifies key kinase regulators and SopB-dependent host phosphorylation events. Sci Signal. 2011. doi:10.1126/scisignal.2001668

24. Cheng S, Wang L, Liu Q, Qi L, Yu K, Wang Z, et al. Identification of a Novel *Salmonella* Type III Effector by Quantitative Secretome Profiling. Mol Cell Proteomics. 2017. doi:10.1074/mcp.RA117.000230

25. Qi L, Hu M, Fu J, Liu Y, Wu M, Yu K, et al. Quantitative proteomic analysis of host epithelial cells infected by *Salmonella enterica* serovar Typhimurium. Proteomics. 2017. doi:10.1002/pmic.201700092

26. Noster J, Chao TC, Sander N, Schulte M, Reuter T, Hansmeier N, et al. Proteomics of intracellular *Salmonella enterica* reveals roles of *Salmonella* pathogenicity island 2 in metabolism and antioxidant defense. PLoS Pathog. 2019. doi:10.1371/journal.ppat.1007741

27. Reuter T, Vorwerk S, Liss V, Chao TC, Hensel M, Hansmeier N. Proteomic Analysis of *Salmonella*-modified Membranes Reveals Adaptations to Macrophage Hosts. Mol Cell Proteomics. 2020;19: 900–912. doi:10.1074/mcp.RA119.001841

28. D’Costa VM, Coyaud E, Boddy KC, Laurent EMN, St-Germain J, Li T, et al. BioID screen of *Salmonella* type 3 secreted effectors reveals host factors involved in vacuole positioning and stability during infection. Nat Microbiol. 2019. doi:10.1038/s41564-019-0580-9

29. Sontag RL, Nakayasu ES, Brown RN, Niemann GS, Sydor MA, Sanchez O, et al. Identification of Novel Host Interactors of Effectors Secreted by *Salmonella* and Citrobacter. mSystems. 2016;1: 1–12. doi:10.1128/msystems.00032-15

30. Hensel M, Shea JE, Gleeson C, Jones MD, Dalton E, Holden DW. Simultaneous identification of bacterial virulence genes by negative selection. Science (80- ). 1995. doi:10.1126/science.7618105

31. Paetzold S, Lourido S, Raupach B, Zychlinsky A. *Shigella flexneri* phagosomal escape is independent of invasion. Infect Immun. 2007;75: 4826–4830. doi:10.1128/IAI.00454-07

32. Ball B, Sukumaran A, Geddes-McAlister J. Label-free quantitative proteomics workflow for discovery-driven host-pathogen interactions. J Vis Exp. 2020. doi:10.3791/61881 (2020)

33. Ball B, Geddes-McAlister J. Quantitative Proteomic Profiling of *Cryptococcus neoformans*. Curr Protoc Microbiol. 2019;55: e94. doi:10.1002/cpmc.94

34. Wisniewski JR, Gaugaz FZ. Fast and sensitive total protein and peptide assays for proteomic analysis. Anal Chem. 2015;87: 4110–16. doi:10.1021/ac504689z

35. Rappsilber J, Mann M, Ishihama Y. Protocol for micro-purification, enrichment, pre-fractionation and storage of peptides for proteomics using StageTips. Nat Protoc. 2007;2: 1896–1906. doi:10.1038/nprot.2007.261

36. Cox J, Mann M. MaxQuant enables high peptide identification rates, individualized p.p.b.- range mass accuracies and proteome-wide protein quantification. Nat Biotechnol. 2008;26: 1367–1372. doi:10.1038/nbt.1511

37. Cox J, Neuhauser N, Michalski A, Scheltema RA, Olsen J V., Mann M. Andromeda: A peptide search engine integrated into the MaxQuant environment. J Proteome Res. 2011;10: 1794–1805. doi:10.1021/pr101065j

38. Cox J, Hein MY, Luber CA, Paron I, Nagaraj N, Mann M. Accurate Proteome-wide Label-free Quantification by Delayed Normalization and Maximal Peptide Ratio Extraction, Termed MaxLFQ. Mol Cell Proteomics. 2014;13: 2513–2526. doi:10.1074/mcp.M113.031591

39. Tyanova S, Temu T, Sinitcyn P, Carlson A, Hein MY, Geiger T, et al. The Perseus computational platform for comprehensive analysis of (prote)omics data. Nature Methods. 2016. pp. 731–740. doi:10.1038/nmeth.3901

40. Benjamini Y, Hochberg Y. Controlling the False Discovery Rate: A Practical and Powerful Approach to Multiple Testing. J R Stat Soc Ser B. 1995;57: 289–300. doi:10.1111/j.2517-6161.1995.tb02031.x

41. Szklarczyk D, Gable AL, Lyon D, Junge A, Wyder S, Huerta-Cepas J, et al. STRING v11: Protein-protein association networks with increased coverage, supporting functional discovery in genome-wide experimental datasets. Nucleic Acids Res. 2019;47: D607–13. doi:10.1093/nar/gky1131

42. Datsenko KA, Wanner BL. One-step inactivation of chromosomal genes in Escherichia coli K-12 using PCR products. Proc Natl Acad Sci. 2000. doi:10.1073/pnas.120163297

43. Clerc PL, Ryter A, Mounier J, Sansonetti PJ. Plasmid-mediated early killing of eucaryotic cells by *Shigella flexneri* as studied by infection of J774 macrophages. Infect Immun. 1987. doi:10.2105/AJPH.2005.076943

44. 44. Wu J, Pugh R, Laughlin RC, Andrews-Polymenis H, McClelland M, Bäumler AJ, et al. High-throughput Assay to Phenotype *Salmonella enterica* Typhimurium Association, Invasion, and Replication in Macrophages. J Vis Exp. 2014. doi:10.3791/51759

45. Hiraga S, Ichinose C, Niki H, Yamazoe M. Cell cycle-dependent duplication and bidirectional migration of SeqA-associated DNA-protein complexes in *E. Coli*. Mol Cell. 1998. doi:10.1016/S1097-2765(00)80038-6

46. Müller P, Chikkaballi D, Hensel M. Functional dissection of SseF, a membrane-integral effector protein of intracellular *Salmonella enterica*. PLoS One. 2012. doi:10.1371/journal.pone.0035004

47. Hubner NC, Bird AW, Cox J, Splettstoesser B, Bandilla P, Poser I, et al. Quantitative proteomics combined with BAC TransgeneOmics reveals *in vivo* protein interactions. J Cell Biol. 2010. doi:10.1083/jcb.200911091

48. Bustamante VH, Martinez LC, Santana FJ, Knodler LA, Steele-Mortimer O, Puente JL. HilD-mediated transcriptional cross-talk between SPI-1 and SPI-2. Proc Natl Acad Sci. 2008. doi:10.1073/pnas.0801205105

49. Watson KG, Holden DW. Dynamics of growth and dissemination of *Salmonella in vivo*. Cell Microbiol. 2010. doi:10.1111/j.1462-5822.2010.01511.x

50. Osborne SE, Tuinema BR, Mok MCY, Lau PS, Bui NK, Tomljenovic-Berube AM, et al. Characterization of DalS, an ATP-binding Cassette Transporter for d-Alanine, and Its Role in Pathogenesis in *Salmonella enterica*. J Biol Chem . 2012. doi:10.1074/jbc.M112.348227

51. Ball B, Bermas A, Carruthers-Lay D, Geddes-McAlister J. Mass Spectrometry-Based Proteomics of Fungal Pathogenesis, Host–Fungal Interactions, and Antifungal Development. J Fungi. 2019;5: 52. doi:10.3390/jof5020052

52. Retanal C, Ball B, Geddes-McAlister J. Post-Translational Modifications Drive Success and Failure of Fungal–Host Interactions. J Fungi. 2021. doi:10.3390/jof7020124

53. Sukumaran A, Coish J, Yeung J, Muselius B, Gadjeva M, MacNeil A, et al. Decoding communication patterns of the innate immune system by quantitative proteomics. J Leukoc Biol. 2019.

54. Tuinema BR, Reid-Yu SA, Coombes BK. *Salmonella* evades D-amino acid oxidase to promote infection in neutrophils. MBio. 2014;5: 1–9. doi:10.1128/mBio.01886-14

55. Welting TJM, Kikkert BJ, Van Venrooij WJ, Pruijn GJM. Differential association of protein subunits with the human RNase MRP and RNase P complexes. Rna. 2006;12: 1373–1382. doi:10.1261/rna.2293906

56. Lapek JD, Mills RH, Wozniak JM, Campeau A, Fang RH, Wei X, et al. Defining Host Responses during Systemic Bacterial Infection through Construction of a Murine Organ Proteome Atlas. Cell Syst. 2018;6: 579–592.e4. doi:10.1016/j.cels.2018.04.010

57. Pietzner M, Wheeler E, Carrasco-Zanini J, Raffler J, Kerrison ND, Oerton E, et al. Genetic architecture of host proteins involved in SARS-CoV-2 infection. Nat Commun. 2020;11: 1–14. doi:10.1038/s41467-020-19996-z

58. Rasko DA, Sperandio V. Anti-virulence strategies to combat bacteria-mediated disease. Nat Rev Drug Discov. 2010;9: 117–128. doi:10.1038/nrd3013

59. Meissner F, Geddes-McAlister J, Mann M, Bantscheff M. The emerging role of mass spectromerty based proteomics in drug discovery. Nat Rev Drug Discov. 2021;In Press.

60. Mesnage S, Dellarole M, Baxter NJ, Rouget JB, Dimitrov JD, Wang N, et al. Molecular basis for bacterial peptidoglycan recognition by LysM domains. Nat Commun. 2014;5. doi:10.1038/ncomms5269

61. Bastos PAD, Wheeler R, Boneca IG. Uptake, recognition and responses to peptidoglycan in the mammalian host. FEMS Microbiol Rev. 2021;45: 1–25. doi:10.1093/femsre/fuaa044

62. Yokoyama CC, Baldridge MT, Leung DW, Zhao G, Desai C, Liu TC, et al. LysMD3 is a type II membrane protein without an *in vivo* role in the response to a range of pathogens. J Biol Chem. 2018;293: 6022–6038. doi:10.1074/jbc.RA117.001246

63. Willmann R, Lajunen HM, Erbs G, Newman MA, Kolb D, Tsuda K, et al. Arabidopsis lysin-motif proteins LYM1 LYM3 CERK1 mediate bacterial peptidoglycan sensing and immunity to bacterial infection. Proc Natl Acad Sci U S A. 2011;108: 19824–19829. doi:10.1073/pnas.1112862108

